# Endogenous antigen processing promotes mRNA vaccine CD4^+^ T cell responses

**DOI:** 10.1101/2025.03.11.642674

**Authors:** Julia E. Rood, Suh Kyung Yoon, Mary K. Heard, Stephen D. Carro, Emma J. Hedgepeth, Mary E. O’Mara, Michael J. Hogan, Nhu Le, Hiromi Muramatsu, Kieu Lam, Petra Schreiner, Coral Kasden, Hansell H. Stedman, Ryan A. Langlois, James Heyes, Norbert Pardi, Laurence C. Eisenlohr

## Abstract

Lipid nanoparticle (LNP)-encapsulated nucleoside-modified mRNA vaccines elicit robust CD4^+^ T cell responses, which are essential for antiviral immunity^1–3^. While peptides presented to CD4^+^ T cells via major histocompatibility complex class II (MHC II) are traditionally thought to be derived from extracellular sources that are processed by antigen presenting cells (APCs) through the classical exogenous pathway^4,5^, the precise mechanisms of mRNA-LNP vaccine-specific CD4^+^ T cell priming remain unknown. Here, we investigated the role of alternative, endogenous antigen presentation pathways^6,7^ in inducing CD4^+^ T cell responses to mRNA-LNP vaccines. APCs treated with mRNA-LNP vaccines were consistently superior in activating T cells under conditions of endogenous, rather than exogenous, presentation. Immunization with an mRNA-LNP vaccine that excludes antigen expression in APCs resulted in lower antigen-specific CD4^+^ T cell, T follicular helper cell, and antibody responses than mice receiving control vaccine. In contrast, depletion of vaccine antigen from exogenous sources such as muscle cells resulted in little to no reduction in antigen-specific CD4^+^ T cells. Our findings demonstrate that direct presentation of endogenous antigen on MHC II is crucial to mRNA-LNP vaccine-induced immune responses and adds to a growing body of literature that redefines the paradigm of MHC II-restricted antigen processing and presentation.

## INTRODUCTION

In the wake of the COVID-19 pandemic, nucleoside-modified mRNA-lipid nanoparticle (LNP) vaccines have emerged as a transformative technology capable of eliciting targeted, protective immune responses^8^. Their efficacy is largely dependent on the induction of vaccine-specific CD4^+^ T cells, which are critical in orchestrating antiviral defenses^2,3,9,10^. Early CD4^+^ T cell and T follicular helper (Tfh) cell responses elicited by mRNA-LNP vaccination predict later titers of neutralizing antibodies^3,11^, a key correlate of protective immunity^12^.

CD4^+^ T cells become activated upon T cell receptor (TCR)-mediated recognition of cognate peptide presented by major histocompatibility complex class II (MHC II). Convention holds that peptides bound to MHC II derive primarily from extracellular depots of antigen, which are internalized by professional antigen presenting cells (APCs) and processed via the classical “exogenous” pathway^4,5^. In the setting of mRNA-LNP vaccination, the intramuscular route of administration has led many to assume that skeletal muscle cells are a primary source of antigen, given studies that demonstrate high uptake of mRNA-LNP at the injection site^13,14^, and implicate transfer of mRNA-LNP vaccine-derived antigen from myocytes to APCs^15^. Others point to the fact that monocytes, macrophages, and dendritic cells rapidly infiltrate the injection site, avidly take up mRNA-LNP, and translate the encoded antigen, implicating hematopoietic cells as a major source of antigen^13,16–18^. In either case, classical exogenous antigen processing is presumed to account for MHC II-mediated presentation of mRNA-LNP vaccine antigens to CD4^+^ T cells^19–23^.

However, no studies have directly addressed the antigen presentation pathways necessary for mRNA-LNP vaccine-specific CD4^+^ T cell activation. The possibility that alternative, “endogenous” MHC II antigen processing pathways^6,7^, in which antigen synthesized within the APC is directly processed and loaded onto MHC II via non-classical pathways, might play a role has been speculated^24,25^ but not explored. Nor is it fully understood which cells are critical for producing the mRNA-LNP vaccine antigen that a CD4^+^ T cell will ultimately recognize. Extrapolation from antigen expression patterns alone is problematic since high levels of antigen expression do not predict high antigen presenting capacity^26–28^.

We previously showed that in mice infected with influenza virus, the majority of the influenza-specific CD4^+^ T cell response is driven by non-classical presentation of endogenous antigen^29^. Productively infected APCs directly presented endogenously-produced influenza peptides to CD4^+^ T cells, resulting in a far broader and more potent CD4^+^ T cell response than when viral antigen was acquired exogenously – a finding that was independent of dose, co-stimulation, cytokine milieu, activation of pattern recognition receptors, or other intracellular effects of the infection itself^29^. Such endogenous pathways have been implicated in the setting of other viral infections^30,31^ and provide an intriguing possibility in the context of mRNA-LNP vaccines, given their highly efficient transfection of APCs.

Thus, we hypothesized that endogenous presentation of mRNA-LNP vaccine antigen by APCs is critical for driving the vaccine-specific CD4^+^ T cell response. We found that APCs that have taken up mRNA-LNP vaccine and directly present endogenous vaccine-derived antigens *in vitro* are superior at activating T cells. Using microRNA (miR)-directed targeting of mRNA-LNP vaccines in mice, allowing for cell type-specific depletion of mRNA, we found that expression of vaccine antigen within hematopoietic cells is necessary for eliciting optimal CD4^+^ T cell responses to mRNA-LNP vaccines *in vivo*, further supporting a role for endogenous processing. These findings provide mechanistic insight into the immunobiology of mRNA-LNP vaccines that may inform the development of next-generation vaccines and have significant implications for our understanding of MHC II-restricted antigen processing more generally.

## RESULTS

### Endogenous presentation of MHC II-restricted mRNA vaccine antigens *in vitro*

We first investigated the relative capacity of *in vitro*-generated APCs to present mRNA-LNP vaccine-derived antigens endogenously versus exogenously. We created nucleoside-modified, LNP-encapsulated mRNA vaccines encoding either nucleoprotein (NP) or hemagglutinin (HA), two model viral antigens from influenza virus. Each of these proteins contains a CD4^+^ T cell epitope that is restricted by I-A^b^ (the MHC II molecule in C57Bl/6 [B6] mice), can be generated via either conventional exogenous or endogenous processing, and whose presentation can be sensitively quantified using antigen-specific T cell hybridomas; these are referred to as NP-45 and HA-16, respectively^29^. We then treated bone marrow-derived dendritic cells (BMDCs) generated from either B6 mice (I-A^b^) or BALB/c mice (I-A^d^/I-E^d^) with these mRNA-LNPs and, after a washout, co-cultured them with both untouched BMDCs of the alternative mouse strain and the appropriate I-A^b^-restricted T cell hybridoma (Fig. 1a). B6 BMDCs that were treated with NP mRNA-LNPs and subsequently expressed NP endogenously were far more efficient at presenting NP-45 to T cell hybridomas than those that acquired NP exogenously from MHC II-mismatched BALB/c BMDCs (Fig. 1b). Similarly, B6 BMDCs presented HA-16 more efficiently to T cell hybridomas after direct treatment with HA mRNA-LNPs than when co-cultured with HA^+^ BALB/c BMDCs (Fig. 1c). These disparities in antigen presentation could not be accounted for by major differences in antigen load (Extended Data Fig. 1a-c), MHC II expression level (Extended Data Fig. 1d,e), or co-stimulation (which is not required by T cell hybridomas^32^). Of note, while exogenous transfer of antigen among mRNA^+^ BMDCs cannot be excluded, this cannot explain the increased antigen presentation under “endogenous” conditions, as antigen transfer between B6 and BALB/c BMDCs is expected to be similar. We repeated these assays across a range of mRNA-LNP vaccine doses and found that presentation of endogenous vaccine antigen was consistently superior to presentation of antigen acquired exogenously (Fig. 1d) and despite equivalent levels of antigen and MHC II expression (Extended Data Fig. 1f,g).

**Figure 1.**
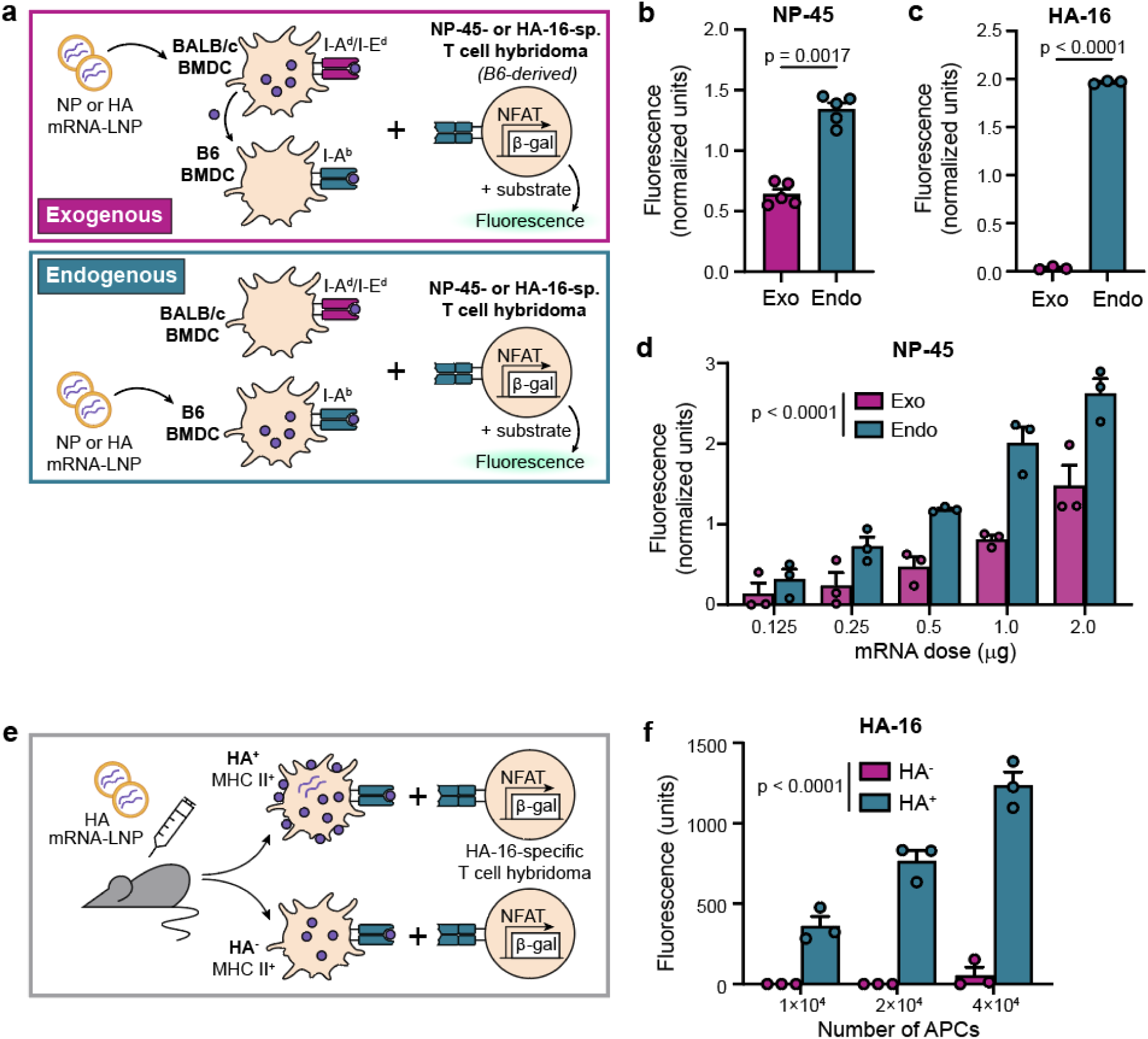
Endogenous presentation of MHC II-restricted mRNA-LNP vaccine antigens is superior to exogenous presentation *in vitro* and *ex vivo*. **a**, Schematic of antigen presentation assay comparing exogenous (“exo”) and endogenous (“endo”) conditions, for panels **b-d**. sp., specific. **b-c**, I-A^b^-restricted presentation of NP-45 peptide **(b)** or HA-16 peptide **(c)**, as measured by fluorescence. Analyzed by paired two-tailed Student’s t-test. **d**, Presentation of NP-45 peptide over a range of mRNA doses. Analyzed by linear mixed-effects model, with pathway (exogenous vs. endogenous) and mRNA dose treated as fixed effects and experiment treated as a random effect; significant main effect of the pathway is shown. In **b-d**, each symbol represents the average fluorescence of technical triplicates over background, normalized to the average fluorescence value within an independent experiment. Bars represent mean +SEM of 5 **(b)** or 3 **(c-d)** independent experiments. **e**, Schematic of *ex vivo* antigen presentation assay in panel **f**. **f**, Presentation of HA-16 peptide by either HA^-^ or HA^+^ primary APCs across a range of APC:T cell hybridoma ratios. Symbols represent technical triplicates. Representative of 2 independent experiments. Analyzed by 2-way ANOVA; significant main effect of the APC type (HA^-^ vs. HA^+^) is shown.

We then sought to determine whether primary murine APCs that had taken up mRNA-LNP vaccine *in vivo* were similarly more efficient at presenting endogenous vaccine antigen. We immunized mice intramuscularly (i.m.) with HA mRNA-LNPs and sorted MHC II^+^ cells with surface expression of HA (HA^+^, directly presenting) or without (HA^-^, indirectly presenting) from the draining lymph nodes (Fig. 1e). HA^+^ cells were confirmed to contain significantly more HA mRNA by quantitative PCR (Extended Data Fig. 1h), consistent with mRNA-LNP vaccine uptake. When cultured with HA-16-specific T cell hybridomas, HA^+^ cells demonstrated far superior presentation of MHC II-restricted HA antigen compared to HA^-^ cells across a range of APC:T cell ratios (Fig. 1f), suggesting that the pool of APCs directly taking up mRNA-LNP vaccine are most capable of activating T cells. The HA^-^ population could contain APCs that acquired HA exogenously as well as those with no exposure to HA and thus may underestimate the effects of exogenous antigen presentation. We also observed nonspecific enhancement of antigen presenting capacity among HA^+^ cells, as they were better at activating T cell hybridomas specific for an irrelevant antigen when pulsed with the cognate peptide (Extended Data Fig. 1i) and expressed slightly higher levels of MHC II compared to HA^-^ cells (Extended Data Fig. 1j). This latter effect is consistent with observations of others^33^ and may be due to the known adjuvant nature of LNPs^34,35^. Nevertheless, these data underscore the finding that APCs taking up mRNA-LNP vaccine and producing vaccine antigen endogenously are considerably more efficient at presenting MHC II-restricted antigen to T cells.

### Validation of mRNA-miR experimental system

We next asked whether endogenous presentation of mRNA-LNP vaccine antigen plays a role in CD4^+^ T cell activation *in vivo*. To investigate this, we added targeting sequences for miR-142, a microRNA (miR) that is expressed exclusively by hematopoietic cells^36^, or a scrambled control to our NP- and HA-encoding mRNA constructs (Fig. 2a). These miR-142-targeted (miR142t) constructs would selectively silence vaccine antigen expression in hematopoietic cells, thus compelling professional APCs to rely on exogenous antigen alone and enabling us to determine the relative contribution of endogenous antigen processing (Fig. 2b). As an additional comparison, we designed NP and HA mRNA constructs with targeting sequences for miR-206 (miR-206t), a skeletal muscle-specific miR^37,38^, to limit one source of exogenous antigen while leaving endogenous antigen processing intact.

**Figure 2.**
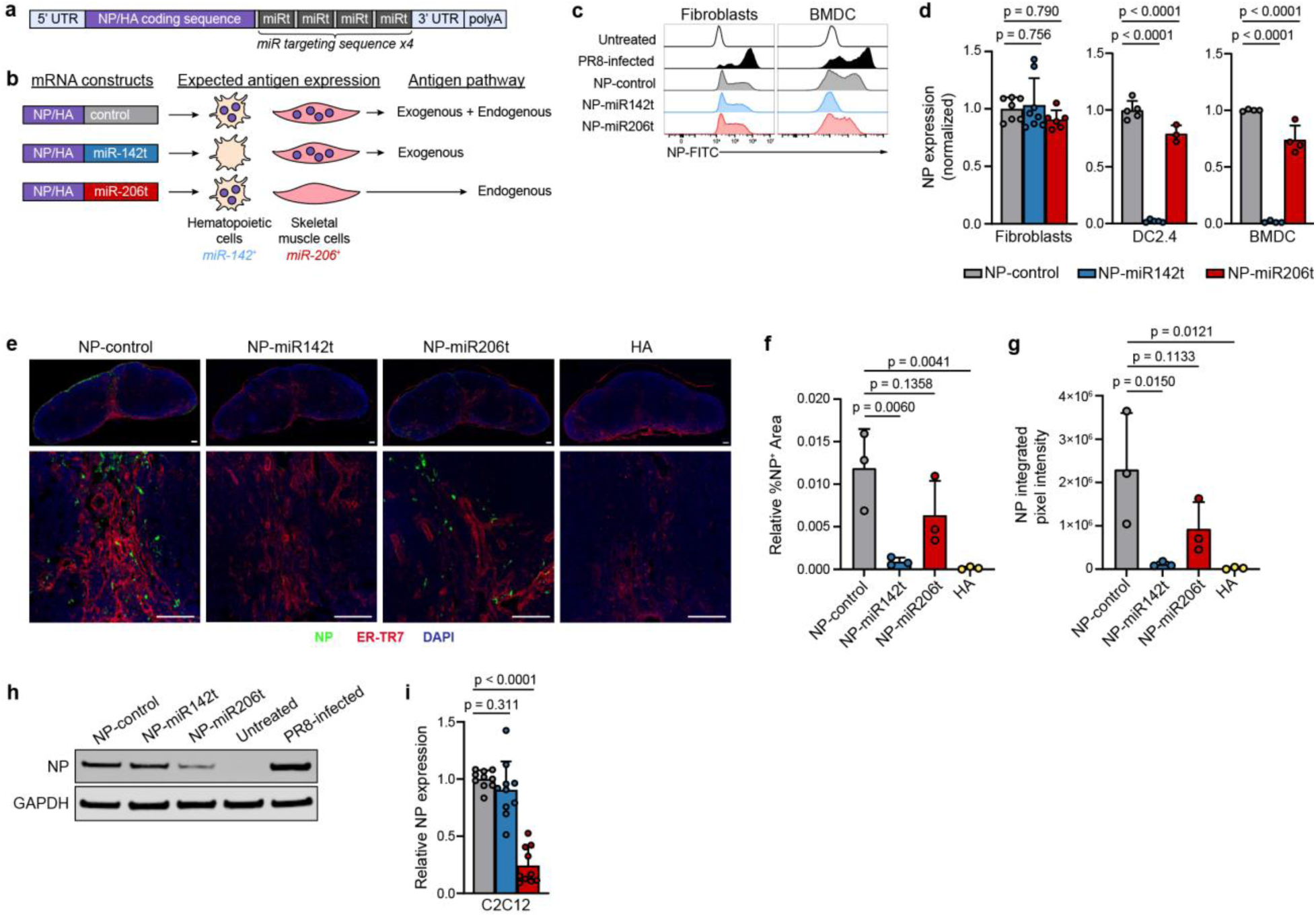
miR targeting enables cell type-specific restriction of mRNA-LNP vaccine antigen. **a**, Schematic of mRNA-miR vaccine constructs encoding influenza NP or HA, followed by miR-targeting sequences in series. **b**, Schematic of expected antigen expression patterns and relevant antigen processing pathways for each mRNA-miR construct. **c**, Representative histograms, gated on live singlets, and **d**, summary data, of NP expression by flow cytometry after *in vitro* transfection with NP-miR mRNA. **e-g**, NP expression in murine lymph nodes after i.m. immunization with mRNA-LNPs. **e**, Representative composite images (top row) and magnified insets (bottom row) of inguinal lymph nodes. ER-TR7 is a fibroblastic reticular cell marker. Scale bars indicate 100 μm. **f**, Surface area of positive NP staining, relative to the entire lymph node surface area. **g**, Integrated intensity of NP staining among all NP^+^ pixels. **f,g**, Bars indicate mean of 3 biological replicates +SEM. Each symbol represents one mouse. Analyzed by 1-way ANOVA; results of Dunnett’s multiple comparison test are shown. **h**, Representative western blot after transfection of C2C12 myotubes with NP-miR mRNA. **i**, Summary data of NP expression as in **h**, quantified relative to GAPDH using densitometry analysis. **d,i**, Symbols represent n=4-8 **(d)** or n=10 **(i)** biological replicates per condition, pooled from 3 **(d)** or 4 **(i)** independent experiments and normalized to the average %NP^+^ value **(d)** or average NP/GAPDH value **(i)** among NP-control-transfected cells, per cell type within each experiment. Analyzed by linear mixed-effects model with cell type **(d)** and mRNA construct **(d,i)** treated as fixed effects and experiment treated as a random effect **(d,i)**. Significance of Dunnett’s multiple comparison test within each cell type is shown. **c,h**, PR8 influenza-infected and untreated controls are included for comparison.

We first validated the specificity of the miR142t constructs by examining differential expression of vaccine-encoded antigen *in vitro*. Transfection of either a dendritic cell line (DC2.4) or BMDCs with hematopoietic-targeted NP-miR142t mRNA completely abrogated NP expression relative to NP-control mRNA, whereas expression was preserved in non-hematopoietic fibroblasts (Fig. 2c,d). To confirm hematopoietic-specific repression of NP expression *in vivo*, NP-miR constructs were encapsulated in LNPs and administered i.m. to B6 mice. Draining lymph nodes were harvested 24 hours later and stained for NP by immunofluorescence. Mice immunized with NP-miR142t mRNA-LNP had significantly lower NP expression, as measured by both surface area and signal intensity of NP staining, compared to mice immunized with NP-control mRNA-LNP (Fig. 1e-g). Of note, a small off-target decrease in NP expression that was observed in hematopoietic cells or tissues exposed to the NP-miR206t construct reached statistical significance *in vitro* but not *in vivo* (Fig. 2d,f,g).

Validation of the miR206t mRNA constructs *in vivo* was challenging due to the known infiltration of inflammatory cells into muscle following immunization; similar to observations from others^39^, many NP^+^ cells localized to the intramuscular connective tissue, confounding measurement of muscle-specific NP expression (Extended Data Fig. 2a). No clear differences were observed in NP expression across muscle tissue sections, as measured by either relative surface area or pixel intensity of positive NP staining (Extended Data Fig. 2b,c), making it difficult to determine the specificity of miR-206 targeting *in vivo*. However, muscle-specific depletion was confirmed by transfection of C2C12 cells, a myoblast cell line that expresses miR-206 upon differentiation to mature myotubes^40^. Differentiated C2C12 cells transfected with NP-miR206t mRNA showed a significant reduction in NP expression compared to cells transfected with either NP-control or NP-miR142t (Fig. 2h,i). These results demonstrate the specificity of the mRNA-miR system, particularly for the hematopoietic-depleting miR142t construct, and allowed us to proceed with examining the impact of each of these constructs on CD4^+^ T cell activation.

### CD4^+^ T cell responses in mRNA-miR-immunized mice

To examine the impact of selectively repressing vaccine mRNA in hematopoietic cells and thus eliminating endogenous processing for MHC II-restricted antigens, we immunized mice i.m. with NP-miR mRNA-LNP vaccines and examined the splenic CD4^+^ T cell response nine days post-immunization (p.i.). The frequency of IFNγ-producing CD4^+^ T cells specific for each of three I-A^b^-restricted epitopes within NP (NP-45, NP-47, NP-52) was substantially decreased in mice immunized with the hematopoietic-depleting NP-miR142t construct compared to NP-control, despite the extreme sensitivity of CD4^+^ T cells^41^ and the protracted opportunity for expansion *in vivo*, consistent with a critical role for endogenous presentation (Fig. 3a). Mice immunized with the muscle-depleting NP-miR206t construct exhibited similar frequencies of NP-specific CD4^+^ T cells compared to controls, suggesting that muscle cells do not constitute a significant exogenous source of antigen for priming vaccine-specific CD4^+^ T cell responses (Fig. 3a). In mice immunized with analogous HA-miR mRNA-LNPs, a similar pattern was observed among HA-16-specific CD4^+^ T cells (Fig. 3b,c). HA-miR142t-immunized mice generated lower frequencies of IFNγ^+^ HA-16-specific CD4^+^ T cells compared to mice immunized with HA-control or HA-miR206t mRNA-LNP, a trend that was most prominent by flow cytometric analysis (Fig. 3c, Extended Data Fig. 3). Mice immunized with HA-miR142t also exhibited a consistent pattern towards lower frequencies of total and TNFα^+^ (but not IL-2^+^) HA-16-specific CD4^+^ T cells (Extended Data Fig. 4a-c). The cytokine profile of HA-16-specific CD4^+^ T cells was similar for all HA-miR mRNA-LNP vaccines (Extended Data Fig. 4d). Overall, these data suggest that the quantity, more than the quality, of vaccine-specific conventional CD4^+^ T cells is dependent on endogenous expression of vaccine antigen within hematopoietic cells.

**Figure 3.**
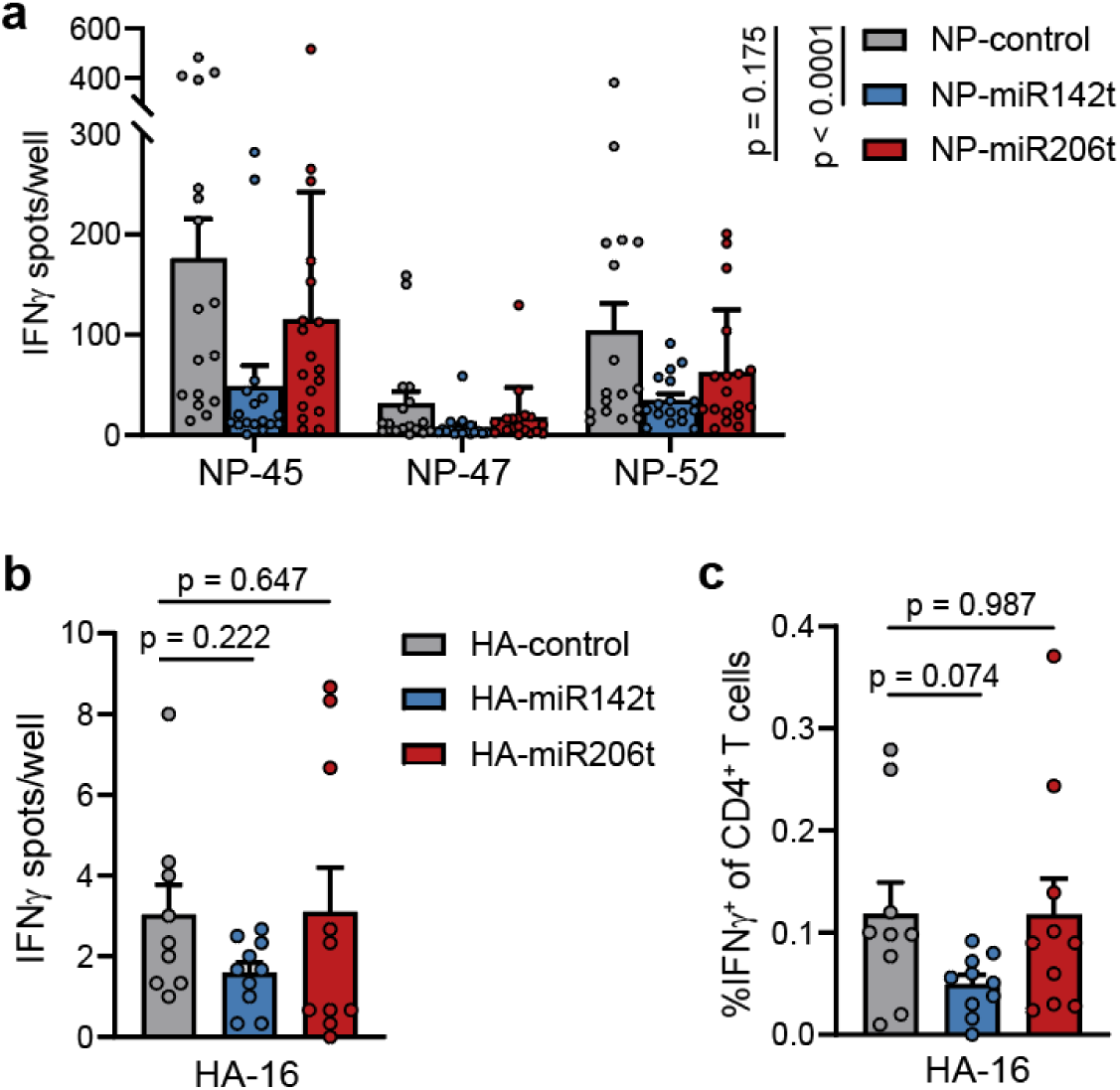
Expression of mRNA-LNP vaccine antigen within hematopoietic cells is required for maximal activation of conventional IFNγ^+^ CD4^+^ T cells. **a**, Frequency of splenic NP-specific CD4^+^ T cells in mice immunized with NP-miR mRNA-LNPs, as quantified by IFNγ ELISpot. Bars represent average of n=17 mice (NP-control, NP-miR142t) or n=18 mice (NP-miR206t) +SEM, from 3 independent experiments. **b-c**, Frequency of splenic HA-16-specific CD4^+^ T cells in mice immunized with HA-miR mRNA-LNPs, as quantified by IFNγ ELISpot **(b)** or IFNγ intracellular cytokine staining **(c)**. Bars represent average of n=9 mice (HA-control) or n=10 mice (HA-miR142t, HA-miR206t) +SEM, from 2 independent experiments. **a-c**, Each symbol represents one mouse. Analyzed by linear mixed-effects model, with mRNA construct modeled as a fixed effect and experiment treated as a random effect. In **(a)**, epitope was also treated as a fixed effect and individual mice were treated as a additional random effect given multiple epitopes tested per mouse. In **(b)**, the number of CD4^+^ T cells plated per well differed between independent experiments so was also treated as a fixed effect. **a-b**, Statistical presented as untransformed values. **a-c**, Significance of pairwise comparisons to control mRNA, corrected for multiple comparisons using Dunnett’s test, are shown.

### Tfh cells and antibody responses in mRNA-miR-immunized mice

Given the essential role of Tfh cells in antibody production and the critical role of antibody in protection against influenza^42^, we next investigated whether restriction of vaccine antigen source, and thus accessible antigen processing pathways, impacted Tfh cell responses. Mice immunized with hematopoietic-depleting HA-miR142t mRNA-LNPs showed a trend towards lower frequencies of Tfh cells and had significantly fewer total Tfh cells in the draining lymph nodes nine days p.i. (Fig. 4a,b and Extended Data Fig. 5), indicating that endogenous antigen expression contributes to Tfh cell activation, similar to its effect on conventional CD4^+^ T cells. Consistent with reduced Tfh cell responses, mice immunized with HA-miR142t also showed lower titers of HA-specific antibodies across multiple timepoints and vaccine doses (Fig. 4c, Extended Data Fig. 6), highlighting the importance of endogenous antigen presentation in shaping the humoral response to mRNA-LNP vaccines. Immunization with muscle-depleting HA-miR206t mRNA-LNPs led to lower numbers of Tfh cells (Fig. 4b), although the effect size was not as large as in HA-miR142t-immunized mice and there were ultimately no differences in antibody titers (Fig. 4c), suggesting that exogenous antigen impacts Tfh cell responses, albeit to a lesser degree.

**Figure 4.**
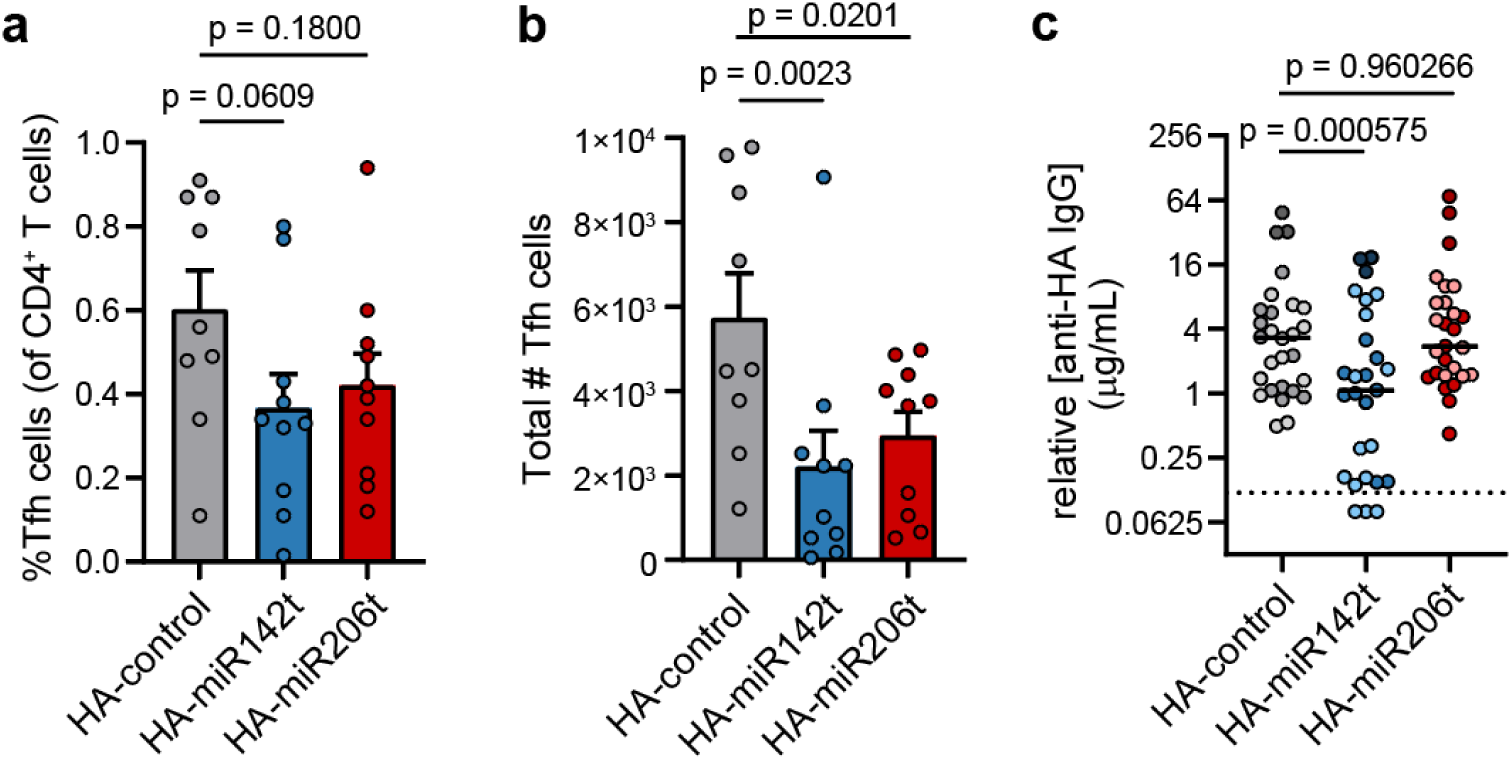
Optimal Tfh cell and vaccine-specific antibody responses require expression of vaccine antigen in hematopoietic cells. **a**, Frequency of Tfh cells, as a percentage of all CD4^+^ T cells, and **b**, total numbers of Tfh cells in the draining lymph nodes of mice immunized with HA-miR mRNA-LNP vaccines. **a-b**, Bars represent average of 9 mice (HA-control) or 10 mice (HA-miR142t, HA-miR206t) +SEM, from 2 independent experiments. Analyzed by linear mixed-effects model, with mRNA construct modeled as a fixed effect and experiment treated as a random effect. Significance of pairwise comparisons to HA-control, determined by Dunnett’s test for multiple comparisons, are indicated. Each symbol represents one mouse. **c**, Serum anti-HA antibody concentration in mice immunized with various doses of HA-miR mRNA-LNPs. Symbols represent individual mouse timepoints, color-coded according to mRNA dose (0.4 μg, light; 4 μg, medium, 8 μg, dark). Data are pooled from 5 independent experiments: 4 μg vaccine (2 experiments, n=10-11 mice/group, day 9 p.i.), 8 μg vaccine (1 experiment, n=3 mice/group, day 9 p.i.), 0.4 μg vaccine (2 experiments, n=7 mice/group, days 14 and 28 p.i.). Solid lines indicate medians. Dotted line indicates lower limit of detection. Analyzed by linear mixed-effects model, with mRNA construct, dose, and timepoint modeled as fixed effects and experiment and individual mice treated as random effects. There was a significant main effect of mRNA construct (p=0.0003516) without interaction with dose (p=0.4023) or timepoint (p=0.88259), allowing for analysis of mRNA-LNP vaccine effects across all timepoints and doses. Significance of pairwise comparisons to HA-control, determined by Dunnett’s test for multiple comparisons, are indicated. Data were log-transformed to meet model assumptions; untransformed data are shown.

## DISCUSSION

Here we examined the role of endogenous MHC II antigen processing and presentation in eliciting CD4^+^ T cell responses to mRNA-LNP vaccines. Contrary to the prevailing paradigm, direct presentation of endogenously-synthesized antigen by APCs is necessary to generate robust vaccine-specific CD4^+^ T cell responses. Repression of vaccine antigen synthesis within APCs leads to fewer conventional CD4^+^ T cells and Tfh cells, as well as lower titers of vaccine-specific antibody. These results provide key insight into the mechanisms underlying induction of effective adaptive immune responses to mRNA-LNP vaccines and highlight the importance of alternative MHC II presentation pathways. Such understanding will inform rational approaches to optimize mRNA-LNP vaccine delivery and design.

APCs are reported to be major sources of mRNA-LNP vaccine antigen *in vivo*^13,16–18^. However, whether antigen-producing APCs are truly essential sources for optimal immune responses has remained unclear, as high antigen and MHC II expression alone do not guarantee effective antigen presentation^26^. Nor was it known whether APCs transfer mRNA-LNP vaccine antigen to other APCs for classical processing or instead directly present their own endogenously expressed antigen. Our study now demonstrates that expression of vaccine antigen within APCs is integral to the downstream adaptive immune response. We show that inhibiting antigen expression in APCs and other hematopoietic cells – crucially, without depleting the cells themselves – significantly reduces the resulting CD4^+^ T cell and Tfh cell responses. This is unlikely to be due to a mere reduction in total body antigen load in miR142t mRNA-LNP-immunized mice, given that when controlling for antigen load *in vitro*, endogenously-produced antigen in APCs is presented far more efficiently than that acquired indirectly from bystander cells. Taken together, our findings support a role for endogenous presentation in driving robust mRNA-LNP vaccine-specific CD4^+^ T cell responses.

Precursor mRNA-LNP vaccine studies observed that dendritic cells transfected with mRNA encoding viral or tumor-associated antigens could directly prime antigen-specific CD4^+^ T cells, in addition to CD8^+^ T cells^43–45^. CD4^+^ T cell responses were initially presumed due to antigen processing through the exogenous pathway^43^, but a later study showed dependence on the proteasome and autophagy^46^, indicating endogenous presentation of cytosolic antigens on MHC II. Numerous other studies have established the ability of viral antigens to be processed via non-classical endogenous MHC II pathways^30,31,47–49^ and to drive antigen-specific CD4^+^ T cell responses to RNA viruses^29^. Thus, it is not unexpected that these pathways could be active in the analogous setting of mRNA-LNP vaccine technology. Antigen may be more abundant within a cell that is actively replicating virus or translating vaccine mRNA into protein, or it may be more readily accessible to endogenous processing pathways. Further characterization of alternative MHC II-restricted antigen processing is needed to better define the intracellular pathways relevant to mRNA-LNP vaccines.

We did not observe a major contribution of skeletal muscle-derived antigen in driving CD4^+^ T cell or antibody responses to intramuscular mRNA-LNP vaccines, although interestingly, mice immunized with the muscle-depleting miR206t construct did generate fewer Tfh cells than controls. Thus, a supplemental role for exogenous antigen presentation cannot be excluded. Given expression of mRNA-LNP vaccine antigen among various populations of stromal cells, such as fibroblasts, endothelial cells, and adipocytes^18^, it seems likely that multiple cell types contribute to CD4^+^ T cell priming through release of antigen and exogenous processing by APCs.

Our study joins a growing body of literature supporting a more nuanced and complex view of MHC II-mediated antigen processing and presentation than the traditional view. Rather than utilizing a single classical exogenous pathway, processing and presentation of MHC II-restricted antigens appears to rely on an interconnected network of intracellular pathways, not unlike direct- and cross-presentation of MHC I-restricted antigens to CD8^+^ T cells^6,7^. The presence of redundancy and complementarity in this system is intuitively appealing and suggests there may be multiple ways to improve protection and durability of responses in future mRNA-LNP vaccines, particularly for immunocompromised individuals or others who mount suboptimal vaccine responses^50^. Future studies are needed to identify the components of endogenous antigen processing pathways that enhance presentation of mRNA-LNP vaccine antigens and target these accordingly in next generation vaccines.

## METHODS

### Cell lines and primary APCs

T cell hybridoma cell lines specific for NP-45 and HA-16 were previously generated in-house^29^ through fusion of splenocytes from PR8-infected B6 mice with BWZ.36/CD8a cells expressing an NFAT-inducible *lacZ* reporter gene^51^. An in-house fibroblast cell line derived from B6 skin fibroblasts and transduced with human class II transactivator (*Ciita)*^52^ was used as a representative non-hematopoietic cell line for *in vitro* studies. These B6-CIITA fibroblasts were maintained in DMEM supplemented with 10% FBS, 2 mM L-glutamine, penicillin, and streptomycin. BMDCs were generated by plating 2×10^6^ murine bone marrow cells (harvested from femurs and tibiae of B6 or BALB/c mice) in 10 mL R10 medium supplemented with 50 ng/mL GM-CSF (VWR 10787-934) in 10 cm tissue culture plates. Medium was supplemented on day 3 and replaced on day 6, prior to BMDC harvest on day 7. BMDCs, DC2.4, and T cell hybridomas were maintained in RPMI supplemented with 10% FBS, 2 mM L-glutamine, 0.05 mM 2-mercaptoethanol, penicillin, and streptomycin (“R10 medium”). C2C12 cells were a kind gift from Foteini Mourkioti (University of Pennsylvania) and maintained in DMEM supplemented with 20% FBS, 2 mM L-glutamine, penicillin, and streptomycin (growth medium). Differentiation into mature myotubes was induced by exchanging growth medium with DMEM containing 2% donor equine serum, 2 mM L-glutamine, penicillin, and streptomycin (differentiation medium)^40^. Differentiation medium was replaced every 24 hours up to the 72-hour timepoint, after which media was changed every 12 hours until harvest, as previously described^53^. T cell hybridomas and any adherent cells transfected with HA mRNA or treated with HA mRNA-LNPs were detached from plates using calcium/magnesium-free medium (made in-house: deionized water with 140 mM NaCl, 2.7 mM KCl, 8.1 mM HNa_2_PO_4_ 7H_2_O, 1.5 mM KH_2_PO_4_, 0.8 mM Na_2_EDTA, phenol red) to avoid cleavage of surface HA; all other adherent cells were detached using trypsin.

### Viruses and infections

Influenza A/Puerto Rico/8/1934 (PR8) was used as a positive control for NP and HA expression *in vitro*. Mouse lung-adapted PR8 was originally provided by Carolina López (Washington University) and grown, harvested, and titered as previously described^54^. B6-CIITA, DC2.4, BMDCs, or C2C12 cells were infected by resuspending in 1 mL PBS containing 1% bovine serum albumin (BSA) and 50 hemagglutinating units (HAU) PR8 and incubating at 37°C with gentle agitation every 15 minutes. After 45 minutes, cells were washed in R10 medium, plated in 6-well plates, and incubated 16-20 hours at 37°C alongside mRNA-transfected experimental samples.

### Synthetic peptides

The following synthetic peptides were used: NP_264–280_ (LILRGSVAHKSCLPACV, “NP-45”), NP_276-292_ (LPACVYGPAVASGYDFE, “NP-47”), NP_306–322_ (LLQTSQVYSLIRPNENP, “NP-52”), and HA_91-107_ (RSWSYIVETPNSENGIC, “HA-16”). All peptides were obtained at >85% purity from GenScript, reconstituted in DMSO, and stored at -20°C.

### Recombinant proteins

Recombinant PR8 HA protein was produced in-house in ExpiCHO-S cells (Gibco A29127) using a PR8 HA plasmid kindly provided by Scott Hensley (University of Pennsylvania). HA protein was purified using HisPur Ni-NTA resin (Thermo Scientific 88221) and concentrated in Amicon 10 kDa MWCO Ultra Centrifugal Filters (Millipore UFC8010).

### mRNA-LNP vaccines

NP-miR and HA-miR sequences were generated using coding sequences for PR8 NP (GenBank accession no. AF389119.1) or HA (GenBank accession no. AF389118.1; ref 55) that were codon-optimized (DNAStar SeqBuilder), followed by four copies of the target sequences for miR-142-3p, miR-206-3p, or a scrambled control, as previously described^56^. These constructs were cloned into an mRNA production plasmid, and sequencing was confirmed through the University of Pennsylvania Genomics and Sequencing Core (RRID:SCR_024999). Nucleoside-modified mRNA was *in vitro* transcribed using the MEGAscript T7 Transcription Kit (Invitrogen AMB13345) and N1-methylpseudouridine (TriLink N-1081), co-transcriptionally capped using CleanCap (TriLink N-7413), and purified by adsorption to cellulose (Sigma-Aldrich 11363), as previously described^57^. mRNA length and integrity were confirmed by agarose gel electrophoresis. Lipid nanoparticle encapsulation using LNP64 was performed by Genevant Sciences Corporation. mRNA-LNP vaccines were stored at -80°C.

### Mice and immunizations

C57Bl/6 (B6) and BALB/c mice were purchased from The Jackson Laboratory and housed in a specific pathogen-free facility at Children’s Hospital of Philadelphia (CHOP). Seven- to nine-week-old male and female mice were immunized by intramuscular injection of mRNA-LNPs diluted in PBS to a total volume of 20 μL in the gastrocnemius muscle. Unless otherwise indicated, a dose of 4 μg mRNA-LNPs was used. All animal studies were performed with the approval of the CHOP Institutional Animal Care and Use Committee.

### Tissue isolation for *in vitro* analysis and flow cytometry

Spleens and draining lymph nodes (inguinal and popliteal) were harvested into RPMI, mechanically disrupted through 70 μm cell strainers using a syringe plunger, ACK lysed for 2 minutes (splenocytes only), re-filtered through 70 μm strainers (splenocytes only), washed, and resuspended in R10 medium to generate single-cell suspensions. Blood was collected from the submandibular vein (anesthetized mice) or by cardiac puncture (immediately after euthanasia) into Serum Gel CAT micro sample tubes (Sarstedt 41.1378.005) and spun at 10,000 × g for 10 minutes. Serum was collected and stored at -20°C.

### Flow cytometry and antibodies

Single-cell suspensions were first stained with LIVE/DEAD Fixable Dead Cell Stain Kits in Aqua (Invitrogen L34957) or Near-IR for 633 or 635 nm excitation (Invitrogen L34975) to exclude dead cells, then were incubated with 10 μg/mL anti-CD16/CD32 (BioXCell BE0307) for Fc receptor blockade. Staining with fluorochrome-conjugated antibodies was performed for 30 minutes at 4°C, unless otherwise indicated. Staining with CXCR5-biotin antibody (SPRCL5) was performed for 60 minutes at 4°C, prior to secondary stain with streptavidin-BV421. All flow cytometry antibodies used are listed in Extended Data Table 2. Monoclonal Anti-Influenza A Virus HA antibody (clone IC5-4F8) was obtained through BEI Resources, NIAID, NIH (NR-48783) and conjugated to AF488 using the Alexa Fluor Antibody Labeling Kit (Invitrogen A20181) according to manufacturer instructions. For intracellular cytokine staining, cells were fixed and permeabilized using the BD Cytofix/Cytoperm Fixation/Permeabilization kit (BD Biosciences 554714). For transcription factor staining, cells were fixed and permeabilized using the eBioscience Foxp3/Transcription Factor Staining Buffer Set (Invitrogen 00-5523-00). Intracellular NP staining was performed after fixation and permeabilization using BD Cytofix/Cytoperm and was not markedly different compared to cells fixed and permeabilized using the eBioscience kit (data not shown). Samples were compensated with UltraComp eBeads (Invitrogen 01-2222-42) and ArC Amine Reactive Beads (Invitrogen A10346) and acquired on CytoFLEX S (3 laser) or CytoFLEX LX cytometers (Beckman Coulter) in the CHOP Flow Cytometry Core Laboratory (RRID:SCR_009726). Data were analyzed in FlowJo v10.8.1 (BD).

### *In vitro* antigen presentation assay

5×10^5^ B6 or BALB/c BMDCs were plated in in 6-well plates and either left untreated or were treated with vaccine by adding mRNA-LNPs in a dropwise fashion. Unless otherwise indicated, 1 μg mRNA-LNPs was used per 5×10^5^ cells. BMDCs were incubated at 37°C for 16-20 hours, then harvested and washed twice to remove residual mRNA-LNPs. A subset of these BMDCs were stained with LIVE/DEAD, Fc blocked, surface-stained with anti-I-A^b^ and anti-I-E^d/k^ antibodies, fixed/permeabilized, and stained with anti-NP or anti-HA antibody to enable comparison of antigen load and MHC II expression. The remaining BMDCs were co-cultured with NP-45- or HA-16-specific T cell hybridomas at a 1:1:2 ratio (5×10^4^ BALB/c BMDCs, 5×10^4^ B6 BMDCs, 1×10^5^ T cell hybridomas per well) in R10 medium in black 96-well plates (Corning 3916) at 37°C for 16-20 hours. For “exogenous” conditions, mRNA-LNP-treated BALB/c BMDCs were combined with untreated B6 BMDCs; for “endogenous” conditions, untreated BALB/c BMDCs were combined with mRNA-LNP-treated B6 BMDCs. NP-45 or HA-16 peptide was added to positive control wells at a final concentration of 10 μg/mL; an equivalent volume of DMSO diluted in R10 medium was added to all other wells. Hybridoma activation was measured by detection of fluorometric β-galactosidase substrate 4-methylumbelliferyl-β-D-galactopyranoside (MUG; Sigma-Aldrich M1633) as previously described^58^. Unless otherwise indicated, background fluorescence from the negative control (untreated BALB/c and B6 BMDCs co-cultured with T cell hybridomas) was subtracted from that of experimental samples. This background level was similar to that of untreated BALB/c BMDCs co-cultured with HA mRNA-LNP-treated B6 BMDCs and NP-45-specific T cell hybridomas.

### Ex vivo antigen presentation assay

Forty-eight hours post-immunization with HA mRNA-LNPs, inguinal and popliteal lymph nodes ipsilateral to the site of intramuscular injection were harvested into R10 medium, pierced with the tip of a 25 G needle to disrupt the capsule, and enzymatically digested with 20 μg/mL DNase I (Roche 10104159001) and 1 mg/mL collagenase IV (Gibco 17104019) in PBS supplemented with 2% FBS for 30 minutes at 37°C. Lymph nodes were then mechanically disrupted through 70 μm cell strainers to generate a single-cell suspension and enriched for MHC II^+^ cells by positive selection with anti-MHC Class II MicroBeads (Miltenyi Biotec 130-052-401). Cells were then stained with LIVE/DEAD Near-IR, Fc blocked with anti-CD16/CD32, and surface-stained with MHC Class II I-A^b^ Monoclonal Antibody (AF6-120.1) conjugated to APC and anti-HA (IC5-4F8) conjugated to AF488. Surface HA^+^ MHC II^+^ and HA^-^ MHC II^+^ APCs were isolated by fluorescence-activated cell sorting on a FACSAria II (BD). A subset of these APCs were analyzed for HA mRNA by qPCR, described below. For the remainder, doubling serial dilutions of APCs (starting at 4×10^4^) were co-cultured with a constant 1.6×10^4^ HA-16-specific T cell hybridomas per well in black 384-well plates (Corning CLS3571) at 37°C. After 18 hours, hybridoma activation was measured via MUG substrate, as above. MHC II^+^ lymph node cells isolated by FACS from an NP-mRNA-LNP-immunized mouse and plated with HA-16-specific T cell hybridomas provided a background level of fluorescence for each APC:T ratio, which was subtracted from values obtained with HA^+^ and HA^-^ APCs. As a control for general antigen presentation capacity, HA^+^ and HA^-^ APCs were also co-cultured with NP-45-specific T cell hybridomas in the presence of 10 μg/mL NP-45 peptide.

### RNA isolation and RT-qPCR

Total RNA was extracted from sorted HA^+^ and HA^-^ APCs using the RNeasy Mini Kit (Qiagen 74104) and reverse-transcribed to cDNA using the SuperScript III First-Strand Synthesis System (Invitrogen 18080051). Quantitative PCR was performed using the PowerUP SYBR Green PCR Master Mix system (Applied Biosystems A25742) on a StepOnePlus Real Time PCR Machine with StepOnePlus software (Applied Biosystems). Expression was quantified relative to the housekeeping gene *Gapdh* using the ΔΔC_T_ method. The following forward (F) and reverse (R) custom primer pairs for PR8 HA were used: ACGAACAAGGTGAACACGGT (F) and ACGTTCGAGTCGTGGAAGTC (R); TACGGGTACCACCACCAGAA (F) and TGTTGAACTCCTTCCCCACG (R). *Gapdh* primers were: TGTCCGTCGTGGATCTGAC (F) and CCTGCTTCACCACCTTCTTG (R).

### Transfection

Cells were transfected in 6-well plates with 1 μg mRNA using *Trans*IT-mRNA Transfection Kit (Mirus Bio, MIR 2250) according to the manufacturer’s instructions, with the following adjustments to reagent volumes: 2.5 μL *Trans*IT-mRNA Reagent and 1.6 μL mRNA Boost Reagent per 1 μg mRNA in a total volume of 130 μL RPMI. Cells were incubated at 37°C for 16-20 hours, then detached and stained for NP expression.

### Western blot

C2C12 cells transfected with NP mRNA were trypsinized and then lysed in RIPA buffer (Cell Signaling Technology 9806S). Lysates were quantified by Pierce BCA Protein Assay (Thermo Scientific 23227) and separated by SDS-PAGE before transfer to nitrocellulose membrane. Membranes were incubated with primary antibodies specific for either GAPDH (Santa Cruz sc-32233, diluted 1:500) or NP (H19-S24 hybridoma supernatant, diluted 1:10) at 4°C overnight. The next day, membranes were incubated at room temperature for 45 minutes with IRDye 800 CW Donkey anti-mouse IgG secondary antibody (LICORbio 926-32212, diluted 1:10,000) and imaged on LICOR Odyssey (LICORbio) or Azure 500 (Azure Biosystems) fluorescent imagers. NP band intensity was quantified on ImageJ by normalization to GAPDH after subtraction of background from both bands.

### Immunofluorescence

Gastrocnemius muscles and inguinal lymph nodes (iLN) were harvested 24 hours after mRNA-miR-LNP immunization. Muscles were immediately frozen in Optimal Cutting Temperature (OCT) medium (Fisher Scientific 23-730-571). iLN were fixed in periodate-lysine-paraformaldehyde (PLP) buffer overnight and equilibrated in 30% sucrose in PBS overnight before embedding in OCT medium and freezing. Cryosections were taken at 16 µm (iLN) or 10 µm (muscle), dried, and stored at -80°C until staining. iLN sections were permeabilized in 0.5% Triton X-100 for 15 minutes, blocked in 5% goat serum in PBS for 30 minutes, and stained overnight with Influenza A NP monoclonal antibody (Clone D67J) conjugated to FITC (Invitrogen MA1-7332, diluted 1:100) and Fibroblast Marker antibody (ER-TR7) conjugated to AF546 (Santa Cruz Biotechnology, sc-73355, diluted 1:50). Muscle sections were fixed in 4% paraformaldehyde, permeabilized and blocked as above, and stained overnight with FITC-conjugated anti-NP antibody (D67J) diluted 1:100, as above. Both iLN and muscle sections were then stained for 1 hour with goat anti-mouse IgG2a AF488 secondary antibody (Invitrogen A21131, diluted to 10 μg) to boost NP signal, followed by staining with DAPI (Abcam AB228549, diluted 1:1000) for 5 minutes. Muscle sections were additionally stained with wheat germ agglutinin (WGA) AF555 (Invitrogen W32464, diluted 1:40) for 10 minutes. All sections were mounted using ProLong Diamond Antifade Mountant (Invitrogen P36970) and imaged on a Leica Stellaris 5 confocal microscope in the University of Pennsylvania Perelman School of Medicine Cell and Developmental Biology Microscopy Core Facility (RRID:SCR_022373). For the DAPI channel, all images were collected following excitation with the 405 nm laser at an intensity of 2% and gain of 5%. For the wheat germ agglutinin channel, images were collected following excitation with the 553 nm laser at an intensity of 2% and gain of 10%. For the ER-TR7 channel, images were collected following excitation with the 553 nm laser at an intensity of 12% and gain of 20%. For the NP channel, images were collected following excitation with the 495 nm laser at an intensity of 12% and gain of 20%. In all instances, images were collected in a 512 x 512 format at speed of 600 with bidirectional imaging. Four line averages were performed and sections were comprised of 10 steps to generate Z sizes ranging from 15-40 μm depending on the tissue section.

### Image analysis

The Fiji distribution of ImageJ2 (https://imagej.net/software/fiji/) was used for all image analyses. Following raw data acquisition, images were processed into maximum intensity projections and subsequently split into individual channels. To generate representative images, for visual clarity, the minimum and maximum values for the NP channel of iLN or muscle sections were adjusted to [9,61] or [9,41], respectively. ER-TR7 was adjusted to [30,255] for all NP-immunized tissue sections and [25,100] for HA-immunized tissue sections. DAPI was adjusted to values ranging from [0,50-142], depending on the relative brightness of the stain. These adjustments were only made for display purposes and all quantitative analysis was performed on raw images, as described below. For iLN and gastrocnemius muscle sections, boundary conditions were established using the ER-TR7 and WGA channels, respectively. In each instance, a 0.5 sigma Gaussian blur (using unscaled units) was applied to soften edges. A threshold that would incorporate all boundaries of the tissue section was then applied. This threshold was converted into a mask, and holes were filled using the “Fill holes” function of Fiji. The “Analyze particle” function was then used to define the region of interest (ROI) as the peripheral boundary of the main tissue section. The ROI area was measured (limited to the threshold) and used as the total area parameter for calculations. For processing the nucleoprotein (NP) channel, the channel was thresholded using defined parameters for each type of tissue section (i.e., [30,255] for lymph node and [25,255] for muscle). This threshold was then analyzed to determine the area and integrated density of the NP signal. The NP area was divided by the total tissue area (determined via ER-TR7 or WGA) to calculate the relative NP area of each tissue section.

### ELISA

Quantification of serum anti-HA antibodies from immunized mice was performed in 96-well MaxiSorp Immuno plates (Thermo Scientific 439454), which were coated overnight with 2 μg/mL recombinant HA protein in PBS at 4°C. Plates were washed and blocked with blocking buffer (PBS with 3% goat serum, 0.5% milk powder, 0.1% Tween 20) for 1 hour with shaking. Two-fold serial dilutions of serum samples were made in blocking buffer, starting at a 1:50 dilution. Control anti-HA monoclonal antibody (H28-D14), a kind gift from S. Hensley (University of Pennsylvania), was also serially diluted in blocking buffer starting at a 0.1 μg/mL dilution. Plates were washed, then diluted sera and control antibody were transferred to ELISA plates and incubated for two hours with shaking at room temperature. Plates were washed, then incubated with goat anti-mouse IgG (H+L) antibody (Southern Biotechnology 1038-05, diluted 1:5,000) for 1 hour with shaking. Plates were washed and 100 μL TMB SureBlue (Seracare 5120-0047) substrate was added to wells for 5 minutes. Wells were quenched with 50 μL of 2M HCl. Absorbance was read at 450 nm. All wash steps were conducted in triplicate using PBS with 0.1% Tween 20.

### Intracellular cytokine staining

1×10^6^ splenocytes per well were stimulated in U-bottom 96-well tissue culture plates in IS10 medium (IMDM supplemented with 10% FBS, 2 mM L-glutamine, 1 mM sodium pyruvate, 0.05 mM 2-mercaptoethanol, penicillin, and streptomycin) in the presence of synthetic peptides (10 μg/mL) and anti-CD28 antibodies (2 μg/mL; Cytek 40-0281). Secretion was inhibited with 5 μg/mL brefeldin A (BioLegend 420601), and cells were incubated at 37 °C for 7 hours. Samples were subsequently stained, fixed, and permeabilized, as above. For each sample, background from an unstimulated control well (DMSO vehicle) was subtracted from the corresponding peptide-stimulated well.

### ELISpot

Splenic CD4^+^ T cells were purified by negative isolation using Dynabeads Untouched Mouse CD4 Cells Kit (Invitrogen 11415D) according to the manufacturer’s protocol. DC2.4 served as APCs and were matured by culturing in R10 medium supplemented with 50 ng/mL recombinant mouse IFNγ (BioLegend 575304) for 24 hours prior to use. 5×10^4^ DC2.4 were co-cultured with 1-2×10^5^ CD4^+^ T cells in MultiScreen-IP Filter Plates (Millipore S2EM004M99) with 10 μg/ml peptide or DMSO vehicle for 18 hours at 37°C. ELISpot assay and plate development were carried out according to manufacturer instructions (BD Biosciences 551881, 557630, 551951). IFNγ spots were imaged and counted on a CTL ImmunoSpot S6 Universal Analyzer.

### Figures

All graphs were generated in GraphPad Prism 10. Figures were composed for publication in Adobe Illustrator. Assay schematics in Figs. 1a, 1e, 2a, 2b are original and were created using Adobe Illustrator. They are modeled in part after images from Biorender.com, which was accessed with a paid subscription that includes permission for journal publishing.

### Statistics

Statistical tests for all data are indicated in the figure legends. Bars represent mean +SEM unless otherwise indicated. Linear mixed-effects models were used to account for variation between independent experiments and/or accommodate missing values. Linear mixed-effects models were conducted using R and lme4^59^, with p values obtained by likelihood ratio tests of the full model with the effect in question against the model without the effect in question. Where relevant, the method of Levy^60^ was used to obtain tests of main effects while modeling an interaction effect. Dunnett’s multiple comparison test was used to compare each experimental group (miR142t, miR206t) to the control. All other data were analyzed in GraphPad Prism 10. QQ plots were visually inspected to ensure normality; in the event of any obvious deviations from normality for data in which exponential expansion was biologically plausible, data were log-transformed and ensured to fit a normal distribution prior to analysis, as indicated in the figure legends.

## Data Availability

Data and reagents that support the findings of this study are available from the corresponding author upon request.

## ACKNOWLEDGMENTS

We thank Edward Behrens for project feedback and statistical guidance, Scott Canna and Audrey John for scientific comments, Em Elliott for guidance on immunofluorescence, Eileen Goodwin for assistance with antibody ELISAs, and Michela Locci and Emily Bettini for Tfh cell staining protocols. This work was supported by the National Institute of Arthritis and Musculoskeletal and Skin Diseases (NIAMS T32-AR076951, J.E.R.); the National Institute of Allergy and Infectious Diseases (NIAID R01AI153064, N.P.); mRNA Pilot Grant (J.E.R., L.C.E.) from the Institute for RNA Innovation of the Perelman School of Medicine at the University of Pennsylvania; Vagelos Undergraduate Research Grant (S.K.Y.) and Grant for Faculty Mentoring Undergraduate Research (S.K.Y., L.C.E.) from the Center for Undergraduate Research & Fellowships at the University of Pennsylvania; and funds from the Children’s Hospital of Philadelphia (J.E.R., L.C.E.).

## AUTHOR CONTRIBUTIONS

J.E.R. conceptualized and oversaw the project, designed and conducted experiments, analyzed and interpreted the data, created the figures, and wrote and edited the manuscript. S.K.Y. designed, conducted, and analyzed *in vitro* validation and antigen presentation experiments. M.K.H. performed tissue sectioning and immunofluorescence studies, conducted western blot and ELISA experiments, and assisted with *ex vivo* T cell studies. S.D.C. assisted with immunofluorescence imaging, performed image analysis, designed qPCR primers, and provided expertise in western blot studies. E.J.H. designed and conducted *ex vivo* antigen presentation assays and provided essential support for animal studies. M.E.O. assisted with mRNA-miR construct design, generated mRNA, and conducted pilot experiments. M.J.H. assisted with mRNA-miR construct design; designed, conducted, and analyzed ICS experiments; and facilitated the collaboration with Genevant Sciences. N.L. assisted with mRNA-miR construct design and generation. K.L., P.S., and J.H. formulated mRNA in LNPs. C.K. and H.H.S. provided expertise for muscle studies. R.A.L. advised on the design of mRNA-miR constructs. H.M. and N.P. contributed mRNA reagents and related expertise. L.C.E. conceptualized and advised the project, interpreted data, and edited the manuscript. All authors reviewed the manuscript.

## EXTENDED DATA

**Extended Data Figure 1.**
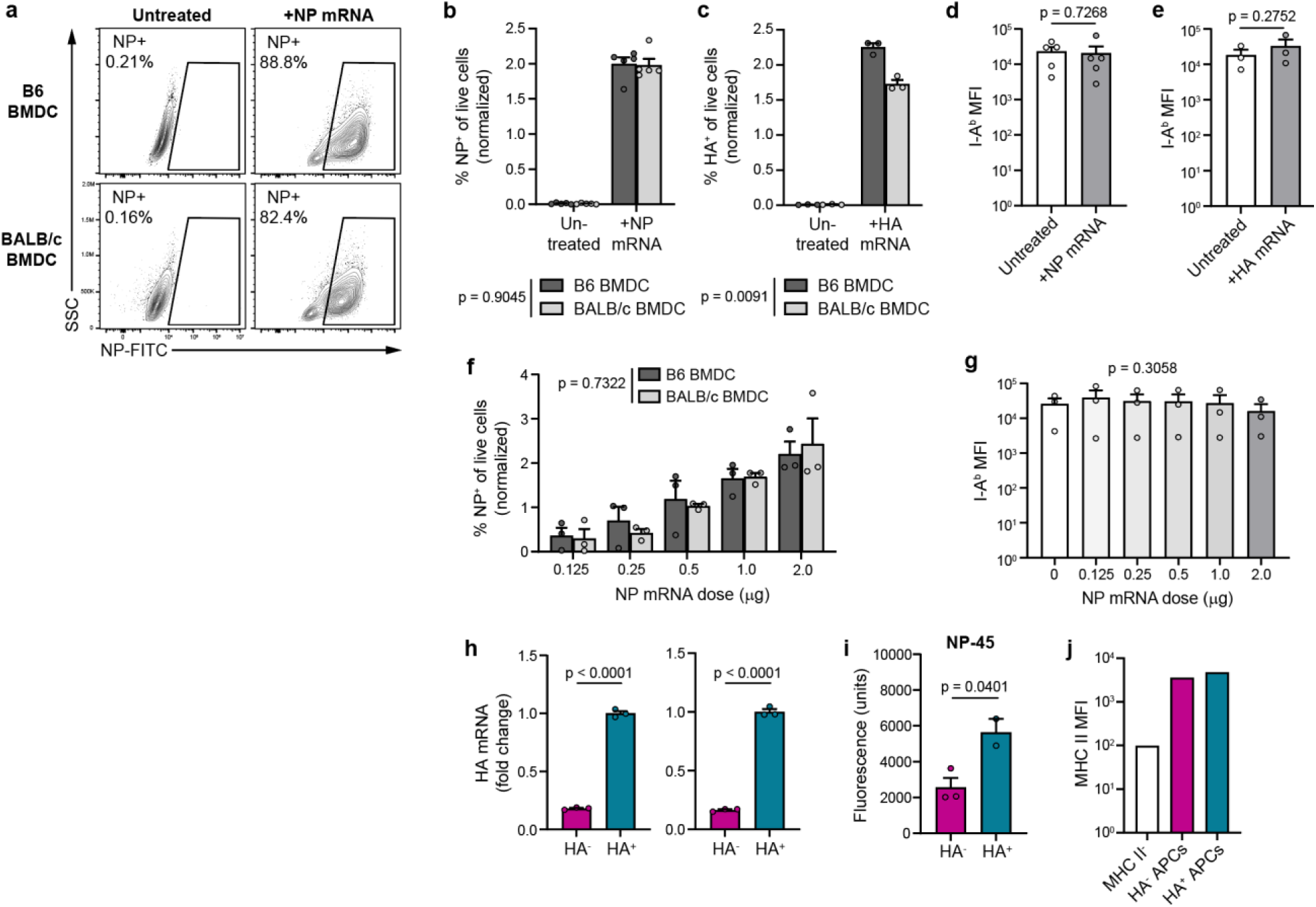
Antigen load and characterization of APCs used in antigen presentation assays. **a**, Representative flow plots of NP staining in BMDCs treated with NP mRNA-LNPs, pre-gated on live singlets. **b**, Summary data indicating %NP^+^ BMDCs, as in **a**. **c**, Percentage of HA^+^ BMDCs after treating with HA mRNA-LNPs, analogous to **b**. **b-c**, Bars represent mean +SEM of n=5 **(b)** or n=3 **(c)** biological replicates per condition, each normalized to the mean value within an independent experiment. Analyzed by 2-way ANOVA; significance of mouse strain main effect is indicated. **d-e**, Median fluorescence intensity (MFI) of I-A^b^ on B6 BMDCs presenting NP **(d)** or HA **(e)** antigen exogenously (untreated) or endogenously (+mRNA). Bars represent mean +SEM of n=5 **(d)** or n=3 **(e)** biological replicates per condition. Analyzed by paired two-tailed Student’s t-presentation assay titration. Bars represent mean +SEM of n=3 biological replicates per condition, each normalized to the mean value within an independent experiment **(f)** or not normalized **(g)**. Data in **f** are analyzed as in **b**. Data in **g** are analyzed by repeated measures 1-way ANOVA. **h**, Relative abundance of HA mRNA by qPCR, using two different primer sets. Bars indicated mean +SEM of technical triplicates. Representative of 3 independent experiments. **i**, Presentation of pulsed NP-45 peptide to T cell hybridomas by sorted HA^+^ and HA^-^ APCs, as measured by fluorescence over background. Bars indicate mean +SEM of technical triplicates (HA^-^) or duplicates (HA^+^). Representative of 3 independent experiments. **j**, MFI of MHC II on sorted HA^-^ and HA^+^ APCs, as well as MHC II^-^ cells for comparison. Representative of 3 independent experiments.

**Extended Data Figure 2.**
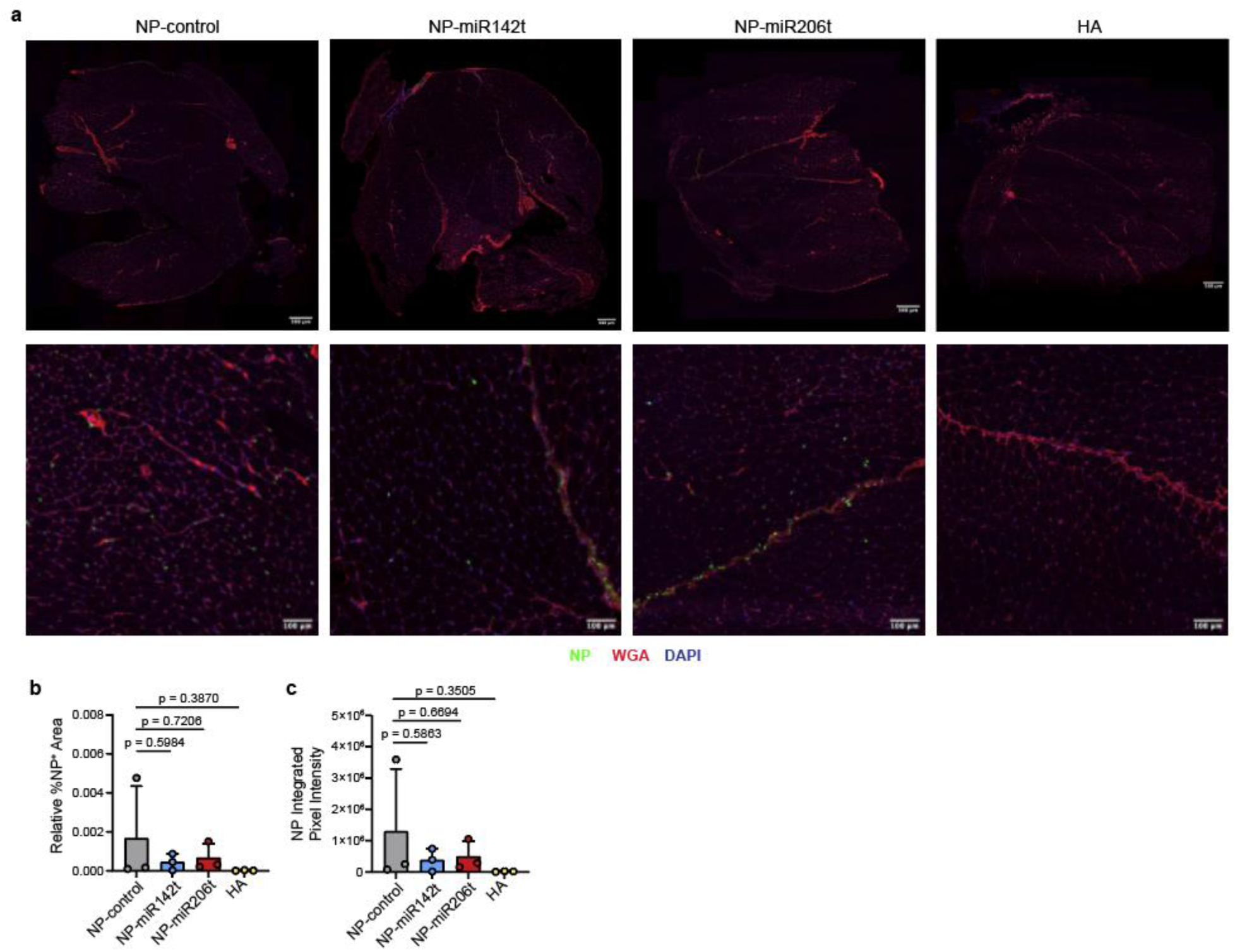
Muscle immunofluorescence analysis of mRNA-miR LNP-immunized mice. **a,** Representative composite images (top row) and magnified insets (bottom row) of gastrocnemius muscle cross-sections. WGA (wheat germ agglutinin) stains muscle cell membranes. **b,** Surface area of positive NP staining, relative to the entire muscle section surface area. **c,** Integrated intensity of NP staining among all NP^+^ pixels. Bars indicate mean of 3 biological replicates +SEM. **b,c,** Analyzed by 1-way ANOVA; results of Dunnett’s multiple comparison test are shown.

**Extended Data Figure 3.**
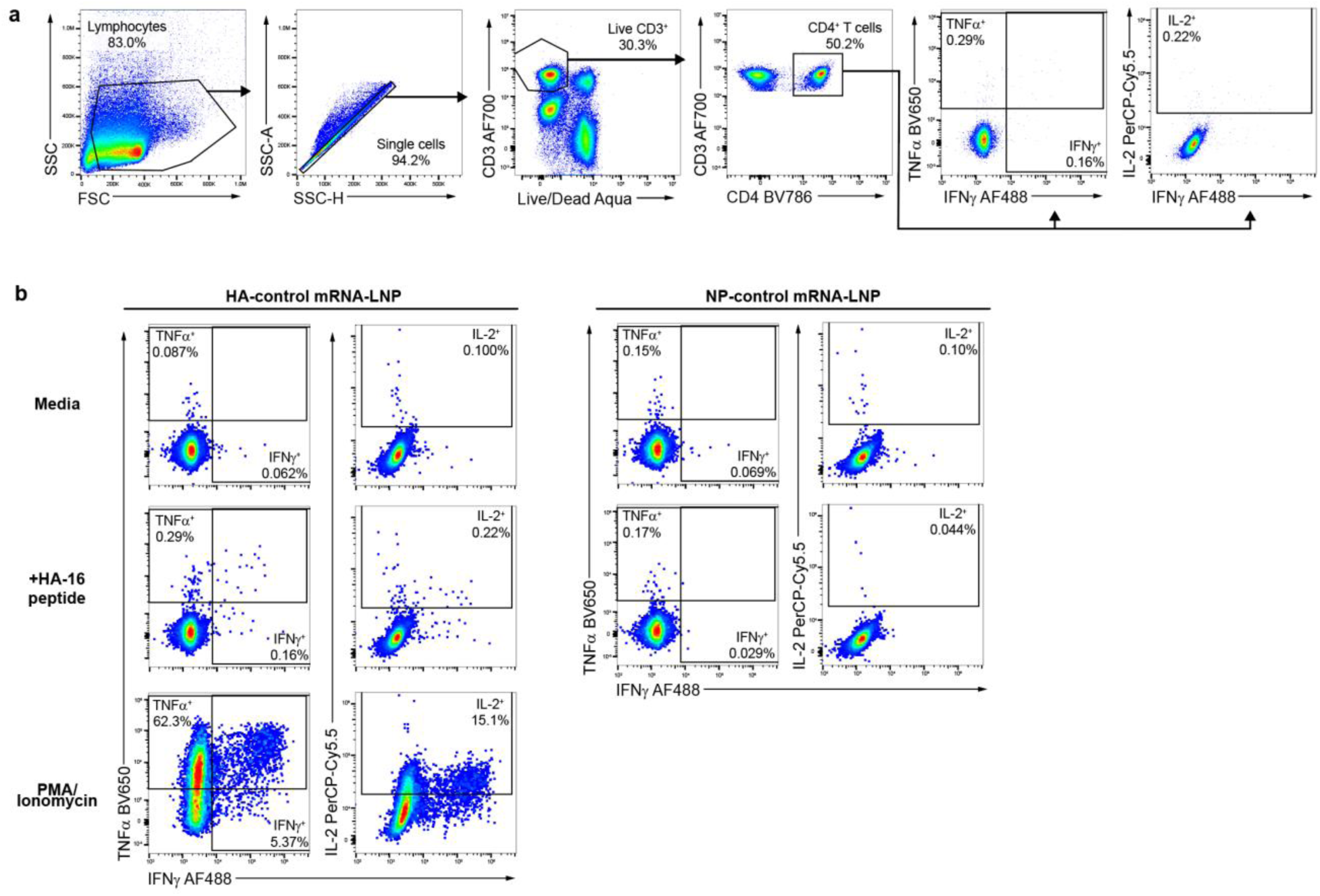
Identification of HA-16-specific CD4^+^ T cells by intracellular cytokine staining. **a**, Gating strategy. Cytokine-producing CD4^+^ T cells were identified as live singlets with a CD3^+^ CD4^+^ phenotype. **b**, Representative flow plots showing CD4^+^ T cell cytokine production from HA-control mRNA-LNP-immunized mice (left panel) at background (top row) and after *in vitro* stimulation with HA-16 peptide (middle row). Negative control (CD4^+^ T cells from an NP-control mRNA-LNP-immunized mouse, right panel) and positive staining control (PMA/Io, bottom row) are also shown.

**Extended Data Figure 4.**
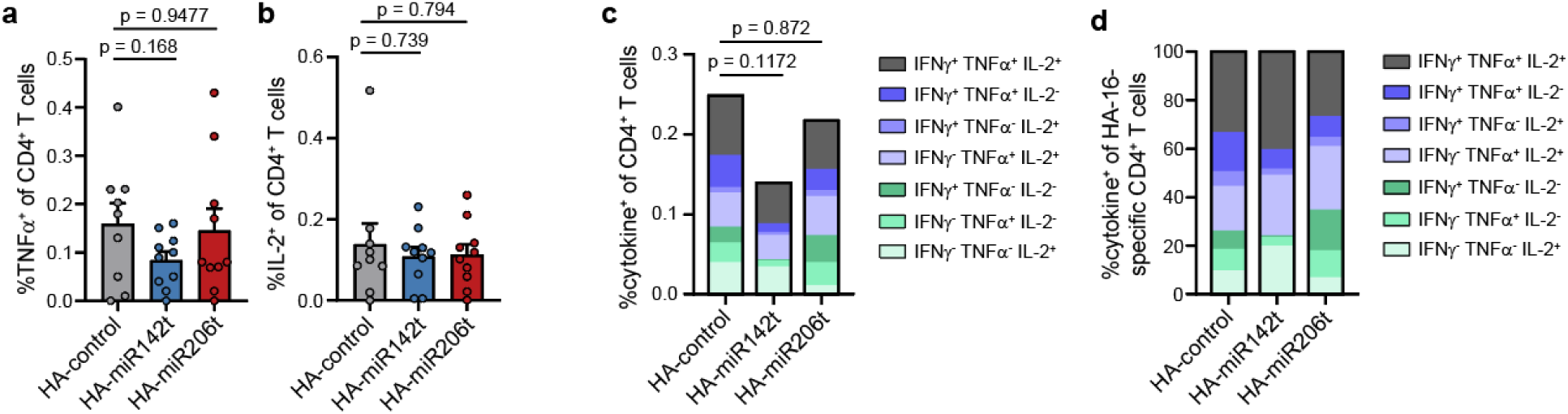
Polyfunctionality of HA-16-specific CD4^+^ T cells. **a-b**, Frequency of splenic HA-16-specific CD4^+^ T cells in mice immunized with HA-miR mRNA-LNP constructs, quantified by intracellular cytokine staining for TNFα **(a)** or IL-2 **(b)**. **c**, Total frequency of HA-16-specific CD4^+^ T cells, defined as specific positivity for IFNγ, TNFα, and/or IL-2. **d**, Proportion of HA-16-specific CD4^+^ T cells producing various subsets of cytokines. **a-d**, Bars represent average of n=9 mice (HA-control) or n=10 mice (HA-miR142t, HA-miR206t) +SEM, from 2 independent experiments. Analyzed by linear mixed-effects model to account for variability between experiments: mRNA construct was modeled as a fixed effect, and experiment was treated as a random effect. In **c** and **d**, individual mice were also treated as a random effect to account for repeated measurements in each mouse. Significance of pairwise comparisons between mRNA groups, indicated in **a-c**, were determined by Dunnett’s multiple comparisons test. In **d**, there was no significant main effect of mRNA construct (p=1.0).

**Extended Data Figure 5.**
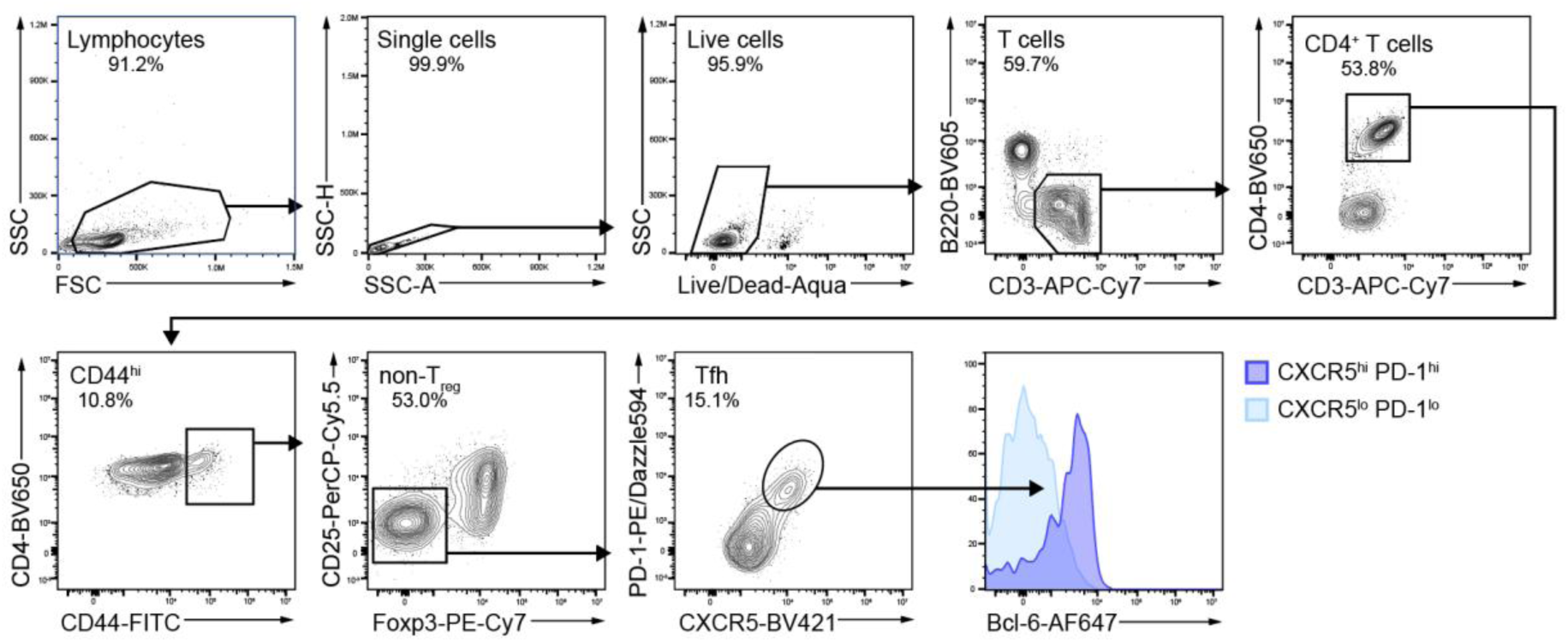
Tfh cell gating strategy. Tfh cells were identified as live singlets with a B220^-^ CD3^+^ CD4^+^ CD44^hi^ Foxp3^-^ CD25^-^ CXCR5^hi^ PD-1^hi^ phenotype. These cells expressed high levels of Bcl-6.

**Extended Data Figure 6.**
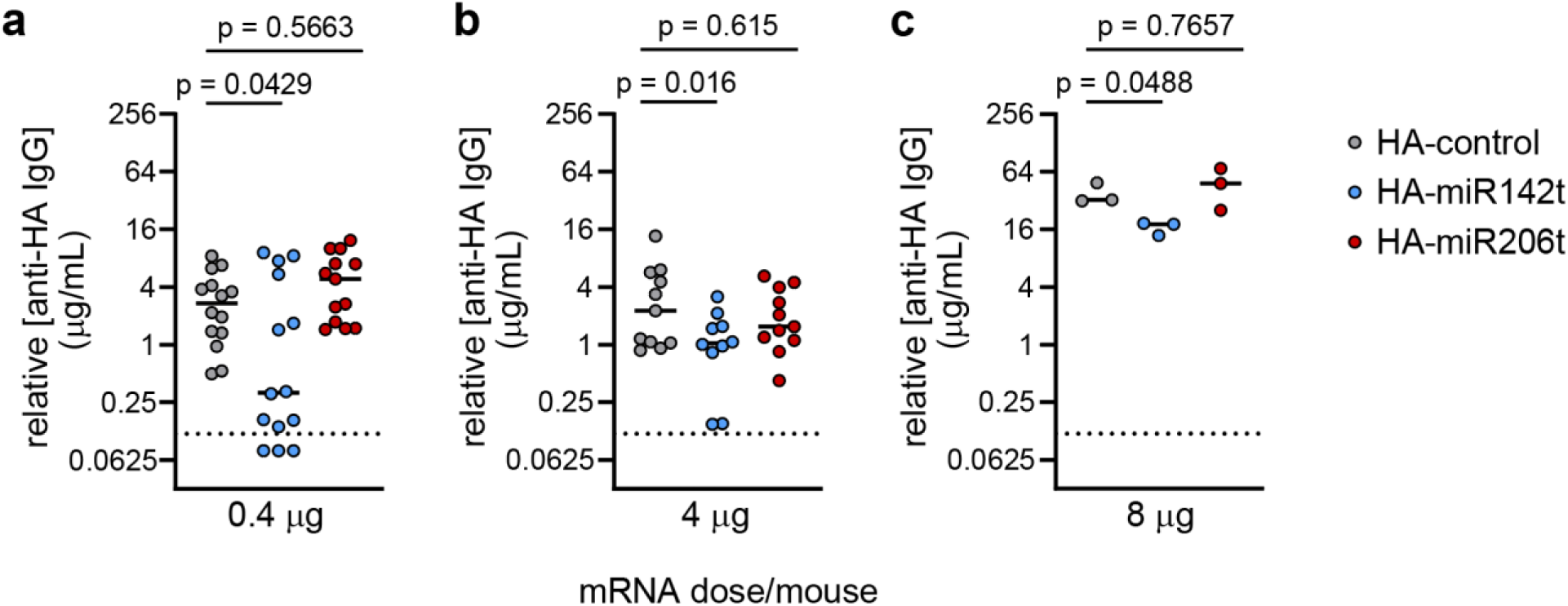
Serum anti-HA antibody concentration by mRNA-LNP vaccine dose. **a-c**, Mice were immunized with 0.4, 4, or 8 μg of HA-miR mRNA-LNPs and serum was analyzed for anti-HA antibodies 9-28 days later. Symbols represent individual mouse timepoints, solid lines indicate medians, and dotted lines indicate lower limit of detection. **a**, Data from n=7 mice/group, each measured at 14 and 28 days p.i., are pooled from 2 independent experiments. **b**, Data from n=11 mice/group (HA-control, HA-miR206t) or n=10 mice/group (HA-miR142t), measured at 9 days p.i., are pooled from 2 independent experiments. **c**, Data from n=3 mice/group, measured at 9 days p.i., from 1 experiment. **a-b**, Analyzed by linear mixed-effects model to account for inter-experiment variability, with mRNA construct treated as a fixed effect and experiment treated as a random effect. In **a**, timepoint is added as a fixed effect and individual mice are added as a random effect to account for sampling the same mice at >1 timepoint. There was a significant main effect of mRNA construct in both models (**a**, p=0.01453; **b**, p=0.03848). **c**, Analyzed by one-way ANOVA (p=0.0270). **a-c**, Significance of pairwise comparisons to HA-control, determined by Dunnett’s test for multiple comparisons, are indicated. Data were log-transformed to meet model assumptions; untransformed data are shown.

**Extended Data Table 1.**
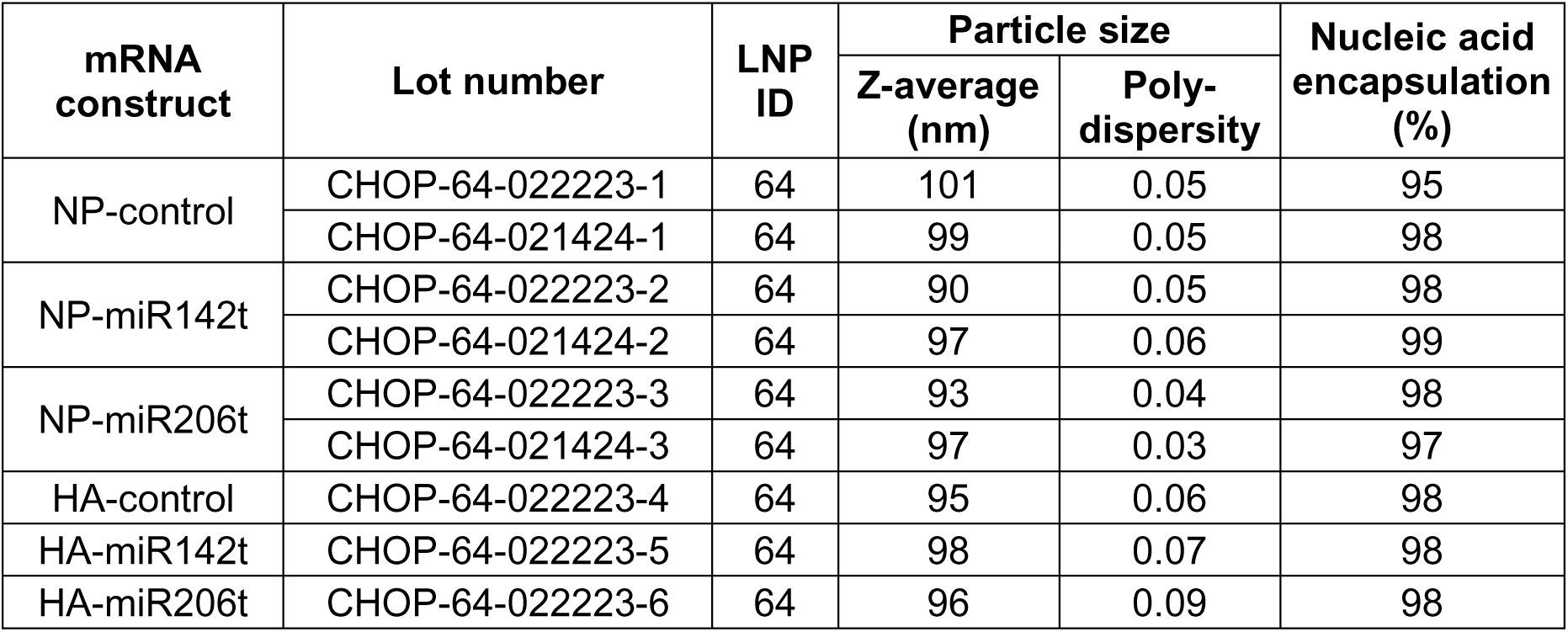
Encapsulation efficiency and particle size of LNPs in mRNA-LNP vaccines.

**Extended Data Table 2.** Antibodies and related staining reagents used for these studies.

| Target | Clone | Supplier | Catalog # | Fluorophore | Lot # |
| --- | --- | --- | --- | --- | --- |
| mouse CD16/CD32 | 2.4G2 | BioXCell | BE0307 | unconjugated | 848823J2 |
| mouse CD185 (CXCR5) | SPRCL5 | Invitrogen | 13-7185-82 | (biotin) | 2691613 |
| (streptavidin) | n/a | BioLegend | 405225 | BV421 | B287245 |
| mouse CD45R/B220 | RA3-6B2 | BioLegend | 103244 | BV605 | B331405 |
| mouse CD4 | RM4-5 | BioLegend | 100545 | BV650 | B417230 |
|  |  |  | 100531 | Pacific Blue | B207592 |
|  |  | BioLegend | 100552 | BV785 | B421544 |
| mouse CD44 | IM7 | BioLegend | 103006 | FITC | B228504 |
| mouse CD25 | PC61.5 | Invitrogen | 45-0251-80 | PerCP-Cy5.5 | 2008275 |
| mouse CD279 (PD-1) | RMP1-30 | BioLegend | 109116 | PE/Dazzle 594 | B278106 |
| mouse CD3e | 145-2C11 | BD Biosciences | 557596 | APC-Cy7 | 7201885 |
|  | 17a2 | BioLegend | 100216 | AF700 | B416228 |
| mouse Foxp3 | FJK-16s | Invitrogen | 25-5773-80 | PE-Cy7 | 2535984 |
| mouse Bcl-6 | K112-91 | BD Biosciences | 561525 | AF647 | 9176164 |
| mouse TNF-α | MP6-XT22 | BioLegend | 506333 | BV650 | B391617 |
| mouse IL-2 | JES6-5H4 | BioLegend | 503822 | PerCP-Cy5.5 | B282323 |
|  |  | BD Biosciences | 554429 | APC | 3051674 |
|  |  | Invitrogen | 12-7021-81 | PE | E030693 |
| mouse IFN-g | XMG1.2 | BioLegend | 505813 | AF488 | B224366 |
| mouse CD8 | 53-6.7 | BioLegend | 100734 | PerCP-Cy5.5 | B408424 |
| mouse I-Ab | AF6-120.1 | Invitrogen | 17-5320-82 | APC | 4289819 |
|  |  | BD Biosciences | 566229 | BV421 | 3163096 |
| mouse I-Ek/Ed/Ep/Er | 14-4-4S | BioLegend | 110211 | AF647 | B331362 |
| influenza A virus HA | IC5-4F8 | BEI Resources | NR-48783 | unconjugated | 70011742 |
| Influenza A virus NP | D67J | Invitrogen | MA1-7332 | FITC | TA2508412 |
|  | IC5-1B7 | BEI Resources | NR-43899 | unconjugated | 70008127 |
| mouse IgG2a | (polyclonal) | Invitrogen | A21131 | AF488 | 2581803 |
| wheat germ agglutinin | n/a | Invitrogen | W32464 | AF555 | 2610352 |
| mouse fibroblast marker | ER-TR7 | Santa Cruz Biotechnology | sc-73355 | AF546 | F1318 |
| mouse IgG (H+L) | (polyclonal) | Southern Biotechnology | 1038-05 | (HRP) | C1717-MK93B |
| mouse GAPDH | 6C5 | Santa Cruz Biotechnology | sc-32233 | unconjugated | K3016 |
| mouse IgG | (polyclonal) | LiCor | 926-32212 | IRDye 800 | D30509-05 |

